# Detecting and controlling dye effects in single-virus fusion experiments

**DOI:** 10.1101/646679

**Authors:** R. J. Rawle, A. M. Villamil Giraldo, S. G. Boxer, P. M. Kasson

## Abstract

Fluorescent dye-dequenching assays provide a powerful and versatile means to monitor membrane fusion events. They have been used in bulk assays, for measuring single events in live cells, and for detailed analysis of fusion kinetics for liposomal, viral, and cellular fusion processes; however, the dyes used also have the potential to perturb membrane fusion. Here, using single-virus measurements of influenza membrane fusion, we show that fluorescent membrane probes can alter both the efficiency and the kinetics of lipid mixing in a dye- and illumination-dependent manner. R18, a dye that is commonly used to monitor lipid mixing between membranes, is particularly prone to these effects, while Texas Red is somewhat less sensitive. R18 further undergoes photoconjugation to viral proteins in an illumination-dependent manner that correlates with its inactivation of viral fusion. These results demonstrate how fluorescent probes can perturb measurements of biological activity and provide both data and a method for determining minimally perturbative measurement conditions.

**Statement of Significance:** Fluorescent dyes are powerful tools for labeling membranes and tracking subcellular objects, and fluorescence dequenching has further been used as a sensitive assay for membrane fusion. Here we show how incorporation of membrane dyes can perturb membrane fusion by influenza virus in a light-dependent manner. We provide a strategy to mitigate this by minimizing dye and light exposure. Finally, we show how in some cases these effects can be due to covalent reaction of some dyes with viral proteins upon illumination. These phenomena may be general and should be carefully controlled for in experiments using such labels.

## Introduction

Fluorescence dequenching has a long history as a means to monitor membrane changes in a variety of biological systems (1-7), but its use has not been without complications. There is a well-documented history of fluorescent probes either not partitioning strongly into the desired “compartment” or exchanging prematurely (8, 9). In early studies of synaptic neurotransmission, there were reports of thermally induced membrane fusion that was ascribed to local heating at high illumination intensity (10). In our prior work, we observed that fusion behavior was more robust with some fluorescent probes than others, and some dyes have been reported by others to perturb lipid-mixing kinetics at high concentrations (3-5 mol%) (3). Both water-soluble and membrane-associated dyes can each perturb fusion; here we examine specifically the effects of membrane-associated dyes that are commonly used to monitor viral membrane fusion. We present a systematic study that compares the effects on influenza virus fusion using two fluorescent probes, R18 and Texas Red, widely used to monitor lipid mixing between a labeled viral particle and a target membrane. Although many other fluorescence-dequenching probes exist, these two were chosen as ones that can readily be introduced into viral membranes at self-quenching concentrations and have previously been reported in single-virus fusion measurements of multiple enveloped viruses (11-14).

The fundamental observation motivating this study is our observation that high illumination intensity can suppress viral fusion yields, as measured by lipid mixing, and such suppression presumably also suppresses downstream stages of fusion. We show that this is dose-dependent in both light intensity and dye concentration and varies with the chemical identity of the dye used. In addition to inhibiting fusion, illumination effects can alter lipid-mixing kinetics. We hypothesize that this is due, at least in part, to photo-induced reactivity between the dye and viral components.

## Materials and Methods

Palmitoyl oleoyl phosphatidylcholine (POPC), Dioleoyl phosphatidylethanolamine (DOPE), and cholesterol (ovine) were purchased from Avanti Polar Lipids (Alabaster, AL). Texas Red-1,2-dihexadecanoyl-sn-glycero-3-phosphoethanolamine (TR-DHPE), Oregon Green-1,2-dihexadecanoyl-sn-glycero-3-phosphoethanolamine (OG-DHPE), and Octadecyl Rhodamine B Chloride (R18) were purchased from Thermo Fisher Scientific (Waltham, MA). Disialoganglioside GD1a from bovine brain (Cer-Glc-Gal(NeuAc)-GalNAc-Gal-NeuAc) was purchased from Sigma-Aldrich (St. Louis, MO). 11-Azidoundecyltrimethoxysilane was obtained from Sikemia (Clapiers, France). 1,1′,1″-tris(1H-1,2,3-triazol-4-yl-1-acetic acid ethyl ester) trimethylamine (TTMA) ligand was a generous gift from Professor Christopher Chidsey at Stanford University. Alkyne-DNA and DNA-lipids were prepared as previously described (15). Ethynyl phosphonic acid was synthesized as previously described (16). Influenza A virus strain X-31 (A/Aichi/68, H3N2) was purchased from Charles River Laboratories (Wilmington, MA).

### Buffer Definitions

Reaction Buffer = 10 mM NaH2PO4, 90 mM sodium citrate, 150 mM NaCl, pH 7.4. Fusion Buffer = 10 mM NaH2PO4, 90 mM sodium citrate, 150 mM NaCl, pH 5.0. HB buffer = 20 mM Hepes, 150 mM NaCl, pH 7.2.

### Microscopy and Illumination Intensity

Fluorescence microscopy images and videos were captured using a Nikon Ti-U epi-fluorescence inverted microscope with a 100x plan apo oil immersion objective, NA = 1.49 (Nikon Instruments, Melville, NY), emission/excitation filter wheels, and an Andor iXon 897 EMCCD camera (Andor Technologies, Belfast, UK). A Spectra-X LED Light Engine (Lumencor, Beaverton, OR) was used as the excitation light source. Metamorph software (Molecular Devices, Sunnyvale, CA) was used to operate the microscope. Filter cubes and excitation/emission filters were as follows: Texas red and R18 images = filter cube (ex = 562/40 nm, bs = 593 nm, em = 624/40 nm), with additional excitation (ex = 560/55 nm) and emission (em = 645/75 nm) filters. Oregon Green images = filter cube (ex = 475/35 nm, bs = 509 nm, em = 528/38 nm), with additional excitation (ex = 460/50 nm) and emission (em = 535/50 nm) filters. All images and video micrographs were captured at 16-bit and 288 ms/frame, with each field of view 82 × 82 µm.

Illumination intensity was measured through the microscope using a handheld optical power meter (Thorlabs, Newton, NJ) capturing all light through a 10x objective, and using the Texas red/R18 filter cube/excitation filters as above. With a 100x objective, this light is distributed across a roughly 50 µm radius circle. From this, we calculated the illumination intensity at the sample surface for the various intensity settings on the LED light engine, with 255 as the max intensity: intensity setting 4/255 = 2.93 W/cm^2^, intensity setting 6/255 = 7.64 W/cm^2^, intensity setting 8/255 = 12.7 W/cm^2^, intensity setting 10/255 = 17.8 W/cm^2^. Prior reports used illumination intensities of 350 W/cm^2^ at 488 nm and 70 W/cm^2^ at 568 nm (17) and approximately 22-28 W/cm^2^ at 532 nm (12).

### Fluorescent labeling of influenza virus membrane

Lipid dye solutions were prepared by mixing either Texas Red-DHPE (0.75 mg/mL in ethanol) or R18 (1.5 mg/mL in ethanol) with HB buffer at the appropriate concentration. A small volume of virus suspension (typically 6 µL) at 2 mg total protein/mL was mixed with 4X volume (typically 24 µL) of dye/HB buffer mixture to yield the final concentration as listed in the data. The solution was incubated for 2 hours at RT on a rocker. Labeled virus was purified from unincorporated dye by adding 1.3 mL HB buffer, and centrifuging for 50 minutes at 21,000 × g. The pellet containing labeled virus was re-suspended in 30 µL of HB buffer, and allowed to rest on ice at 4°C for at least several hours. For lipid mixing assays, labeled virus was stored on ice at 4°C and was used within several days of labeling.

### Estimation of mean dyes per virion

The mean number of dye molecules per virion was estimated by dividing the measured concentration of fluorescent dye in a sample of labeled virus by the estimated concentration of viral particles.

To estimate the mean concentration of fluorescent dyes incorporated into the viral particles, a small volume of labeled virus was solubilized with 10% Triton-X in HB buffer to yield a final Triton-X concentration of 1%. The solubilized virus was diluted in HB buffer with 1% Triton-X and the fluorescence emission spectrum of the sample was measured using a fluorimeter (Perkin Elmer LS 55, Waltham, MA). The integrated fluorescence intensity across the spectrum was calculated and compared to a standard curve of known dye concentration in the same buffer to determine the concentration of fluorescent dye in the viral suspension. Slit widths, excitation wavelengths, and integration windows were optimized for each dye to yield a reasonable signal without saturating the detector. The same settings were then used for all measurements of a given dye. This method approximates the quenching behavior of the dye in the viral membrane, where the effective distribution volume is the membrane volume, as similar to the equivalent concentration of Triton-solubilized dye in solution, using the full solution volume as the distribution volume; deviation from this will contribute to the error of estimation.

Separately, the concentration of virus was estimated as we have reported previously (15) by measuring the viral protein concentration using a BCA protein concentration assay kit (ThermoFisher Scientific) and then dividing the measured protein concentration by the average total protein molecular weight per viral particle (1.8 × 10^8^ Da) calculated from literature values of average copy numbers of viral proteins per virion (18). Mean protein concentration measurements were made on mock labeled virus samples, as the Texas Red and R18 dyes have overlapping absorbance spectrum with the BCA-Cu(I) complex used in the BCA assay.

### Single virus lipid mixing assay

Single virus lipid mixing measurements were performed inside microfluidic flow cells as previously described (15). Briefly, glass coverslips were functionalized with 11-Azidoundecyltrimethoxysilane via vapor deposition. Microfluidic flow cells (2.5 mm × 13 mm × 70 µm) made of poly dimethyl siloxane (PDMS) were then affixed to the functionalized coverslips using epoxy. Subsequently, short DNA oligomers (DNA sequence 5’-alkyne-TCCTGTGTGAAATTGTTATCCGCA-3’) were then attached to the functionalized coverslip surface inside the flow cell via copper catalyzed azide-alkyne click chemistry using TTMA ligand, and the remaining surface was passivated using ethynyl phosphonic acid.

Separately, lipid mixtures containing 67.5 mol% POPC, 20% DOPE, 10% cholesterol, 2% GD1a, 0.5% Oregon Green-DHPE were prepared in chloroform, evaporated to a lipid film under N_2_ gas and house vacuum, resuspended in Reaction Buffer and then extruded into liposomes using a polycarbonate membrane with 100 nm pore size. DNA-lipids (DNA sequence 5’-lipid-TGCGGATAACAATTTCACACAGGA-3’) were incorporated into the outer membrane leaflet of the liposomes at 0.01 mol% by overnight incubation at 4°C. These liposomes displaying DNA were introduced into the microfluidic flow cells. Hybridization between DNA anchored in the liposomes and DNA attached to the functionalized coverslip resulted in DNA-tethered liposomes on the coverslip surface. These tethered liposomes were the target membranes for the lipid assays. Any unbound liposomes were removed by extensive rinsing.

Following preparation of the microfluidic flow cell with target membranes, fluorescently labeled virus was added and incubated with the target membranes for several minutes to allow viral binding. The flow cell was then extensively rinsed with Reaction Buffer to remove any unbound virions. Any images taken to find and focus on a suitable area of the sample during this period were always taken at intensity setting 4/255 on the LED light engine (2.93 W/cm^2^). The buffer in the flow cell was then exchanged with pH 5.0 Fusion Buffer to trigger viral fusion, which was monitored by fluorescence video microscopy at the indicated illumination intensity for five minutes. Lipid mixing events between the viral and target membranes were detected by fluorescence dequenching of the dye-labeled lipid.

### Measurement of lipid mixing in unilluminated areas

To observe lipid mixing in unilluminated areas, microfluidic flow cells with target membranes and bound virus were prepared as above. Before introduction of the pH 5.0 Fusion Buffer, images of bound virus in several areas within the flow cell were taken. Viral fusion was then triggered by introduction of the low pH buffer, during which time none of the areas were illuminated. After approximately five minutes, images of the same areas were again taken. All images were acquired using intensity setting 4/255 on the LED light engine (2.93 W/cm^2^). Lipid mixing events between the viral and target membranes in those areas were also assessed by fluorescence dequenching, as described in the analysis section below. Note that often the same flow cell was used to collect both the data for illuminated and unilluminated areas. In this case, the unilluminated areas were located far away (at least several hundreds of microns) from the illuminated areas to minimize any influence of scattered light on the unilluminated regions during collection of the fusion video.

### Measurement of dye photoadducts

5 µl of labeled virus were applied to an empty microfluidic well on a glass cover slip, illuminated for 1 min at intensity 38/255, and then extracted. Samples were resuspended in 15 µl of loading buffer (50 mM Tris pH 6.8, 2% SDS, 100 mM DTT, 10% glycerol and 0.5% bromophenol blue) and loaded without heating on a 4-12% acrylamide NuPAGE gel (Invitrogen). Proteins were separated through electrophoresis in 50 mM MES, 50 mM Tris Base, 0.1% SDS, 1 mM EDTA, pH 7.3. Fluorescence imaging of the gel (excitation 520-545 nm, emission 577–613 nm) was performed before staining the gel with Coomassie Blue and imaging overall protein content using a ChemiDoc MP instrument (Bio-Rad, Hercules, CA).

### Analysis of microscopy data

Custom-built Matlab programs described previously (15) and available at https://github.com/kassonlab/micrograph-spot-analysis were used to automatically extract fluorescence intensity time traces from individual viral particles in the standard lipid mixing assay, and to analyze those traces to determine the waiting time between pH drop and the onset of lipid mixing.

To estimate the lipid mixing efficiency for the unilluminated data sets, we modified our prior Matlab code to calculate the percentage difference in background subtracted fluorescence intensity per particle in the before and after pH drop images. To determine the appropriate threshold value for the percentage intensity difference which would be categorized as lipid mixing, a control data set was used in which the pH was not dropped, but otherwise all experimental details remained the same. The false positive error in the control data sets were 2% or less.

### Kinetic modeling

Two kinetic models were fit to the single-virus fusion data to test the hypothesis that a dye-driven and light-driven inactivation process could explain the change in fusion efficiencies and rates. Inactivation was assumed to be linear in [dye]・light intensity. Details of the two models follow.

A chemical-kinetics model formalism previously developed for Zika membrane fusion (14) was modified to incorporate three sequential steps from membrane-bound virus to hemifusion in accordance with the single-virus kinetic model used by Floyd and colleagues (11). The resulting formalism had the following kinetic parameters:

B <-> I

B <-> A1 -> A2 -> HF

In the best-fit models, the rate constant for A1->B was zero. These kinetic models were fit to all the observed single-virus fusion data using previously developed code available from https://github.com/kassonlab/zika-kinetics.

A cellular automaton model for influenza virus fusion that tracks activation of individual hemagglutinin trimers was also fit to the single-virus fusion data. This model was original formulated by Ivanovic and co-workers (19, 20) and previously modified by us (21). Code available at https://github.com/kassonlab/ca-fitting was used for nonlinear optimization of rate constants to find rates that maximize the likelihood of the observed single-virus fusion data. Joint fitting was performed across all dye concentrations and light intensities.

## Results and Discussion

To test the effects of fluorescent dye and illumination intensity on influenza membrane fusion, we prepared influenza virus samples labeled with different dye concentrations (see Methods) and performed single-virus fusion experiments in microfluidic flow cells as previously reported (15). In these experiments, labeled influenza virus is bound to GD1a displayed on tethered vesicles (molar ratios 67.5 POPC:20 DOPE:10 Chol:2 GD1a:0.5 OG-DHPE), unbound virus is washed away, and then fusion is triggered by buffer exchange to pH 5.0. Lipid mixing between labeled virus and target membranes is assessed as fluorescence dequenching measured via video microscopy (Fig. 1). Single-event waiting times from pH drop to fluorescence dequenching are compiled into cumulative distribution functions, which are then used to assess both lipid-mixing kinetics and efficiency, which provides an upper bound on fusion efficiency. To assess illumination effects on fusion efficiency, we also measured the fusion efficiency within other fields of view that were illuminated only for two exposures, one prior to triggering fusion and one ∼5 minutes after. These images were used to estimate fusion efficiency for single viral particles and compared to the fusion efficiency observed during illuminated video traces with illumination settings within the range of those reported previously.

**Figure 1:**
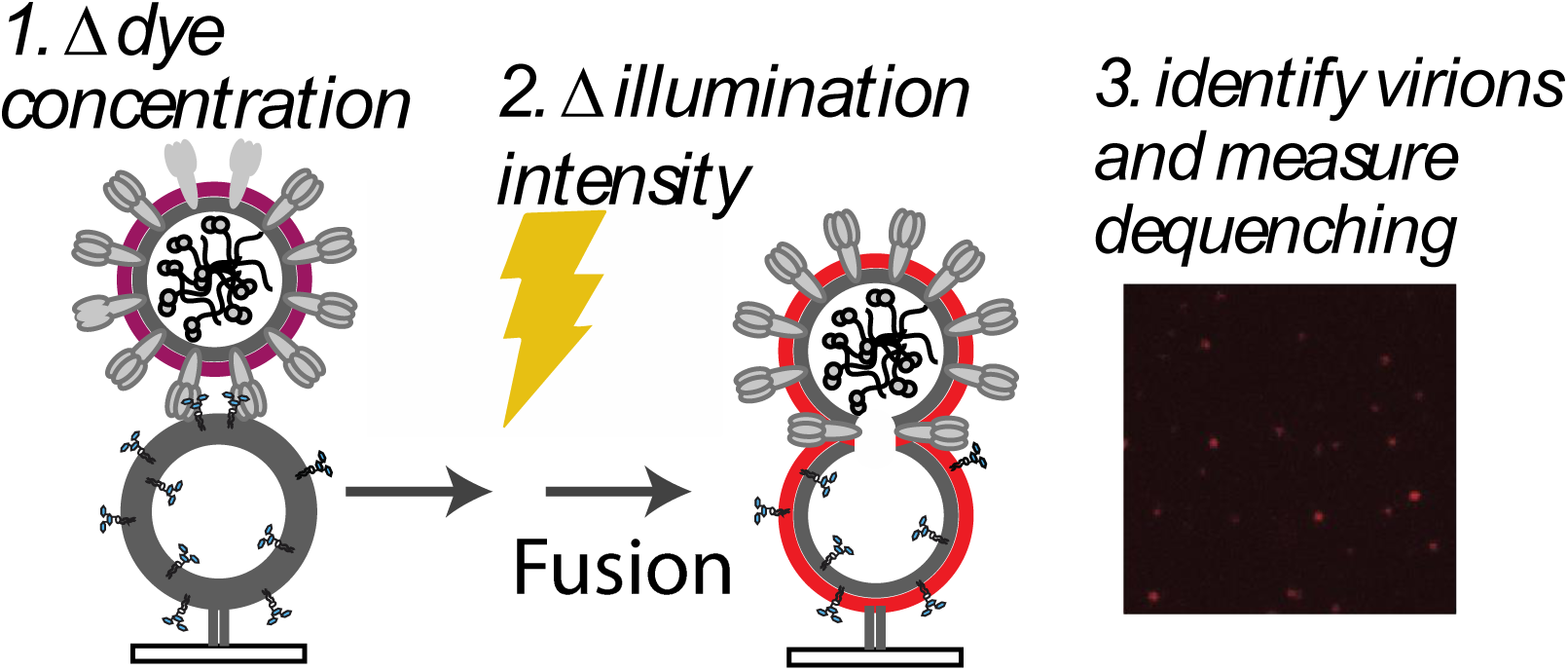
Measurement of single-virus influenza fusion at variable dye concentrations and illumination intensities. X-31 influenza virions are labeled with designated concentrations of Texas Red-DHPE or R18 dye and then bound to GD1a-containing liposomes immobilized inside a microfluidic flow cell. Fusion is triggered by a buffer exchange to pH 5.0, and the sample is subjected to different illumination intensities for 5 minutes. In un-illuminated samples, fusion efficiency is determined by fluorescence dequenching at 5 minutes relative to experiment start; in illuminated samples, lipid-mixing kinetics and efficiency are determined by following time-traces of individual virions identified in the images.

The commonly used R18 dye (octadecylrhodamine B) readily incorporates into influenza viral membranes at self-quenching concentrations (Fig. S1). At the lowest concentration of R18 dye tested and the lowest illumination level, there was no detectable difference in the efficiency of lipid mixing observed from video microscopy traces and that measured in single-time-point experiments. However, a twofold increase in either dye concentration or illumination intensity significantly decreased fusion efficiency, with a more profound effect observed with R18 dye concentration than with illumination (Fig. 2). We also noted changes in fusion efficiency at high R18 concentration independent of illumination consistent with previous reports (3) but no analogous changes with Texas Red (see below); here we concentrate on the illumination-dependent effects. Even at the lowest concentration of dye tested, increasing illumination intensity substantially increased fusion kinetics even as it decreased efficiency, with a ∼2.5-fold variation in t_1/2_ for lipid mixing over a 2.5-fold illumination intensity range (Fig. 3). The intensity distribution of R18-labeled virions changed slightly over the illumination conditions but less than the Texas-Red-labeled virions (Fig. S2), so the decrease in fusion intensity cannot be attributed to detection of poorly-labeled virions at high illumination that then fail to undergo dequenching. These results suggest that some product of dye illumination either a) selectively inactivates slow-fusing viral sub-populations or b) inactivates virus in a time-dependent manner, thus appearing to bias fusion events towards early time-points prior to inactivation.

**Figure 2:**
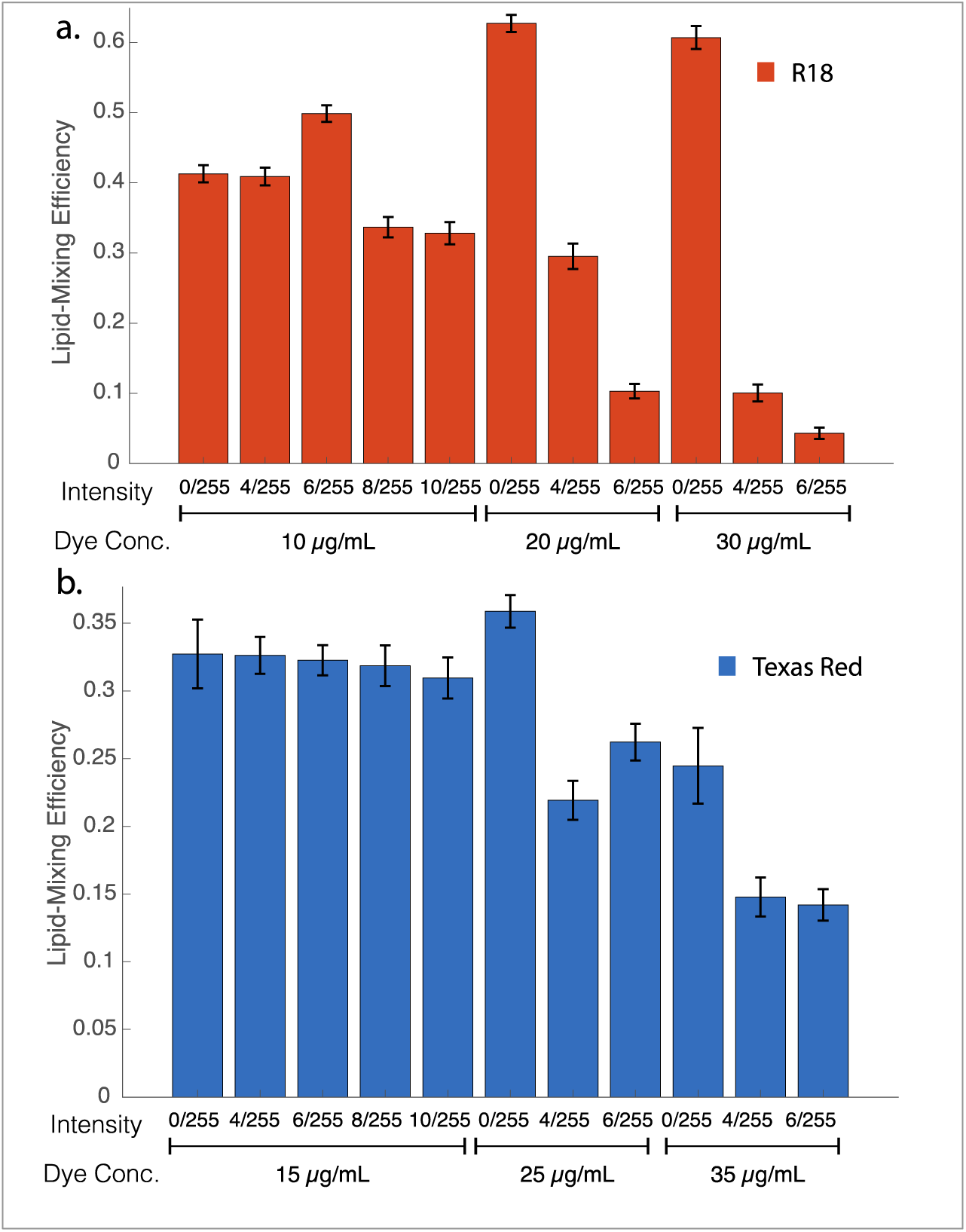
Photoinhibition of membrane fusion by influenza virus labeled with R18 or Texas Red-DHPE dyes. Lipid-mixing efficiency is plotted for virus labeled with different concentrations of (a) R18 or (b) Texas Red dye and exposed to the indicated illumination intensities for 6 min prior to measurement. Photoinhibition is apparent at high dye concentrations and more marked for R18 than Texas Red. Error bars are plotted as standard deviations across fields of view for unilluminated samples and bootstrapped standard deviations across individual viruses for illuminated samples. Illumination intensities are reported as fraction of maximum LED intensity. Measured intensities are given in the Materials and Methods.

**Figure 3:**
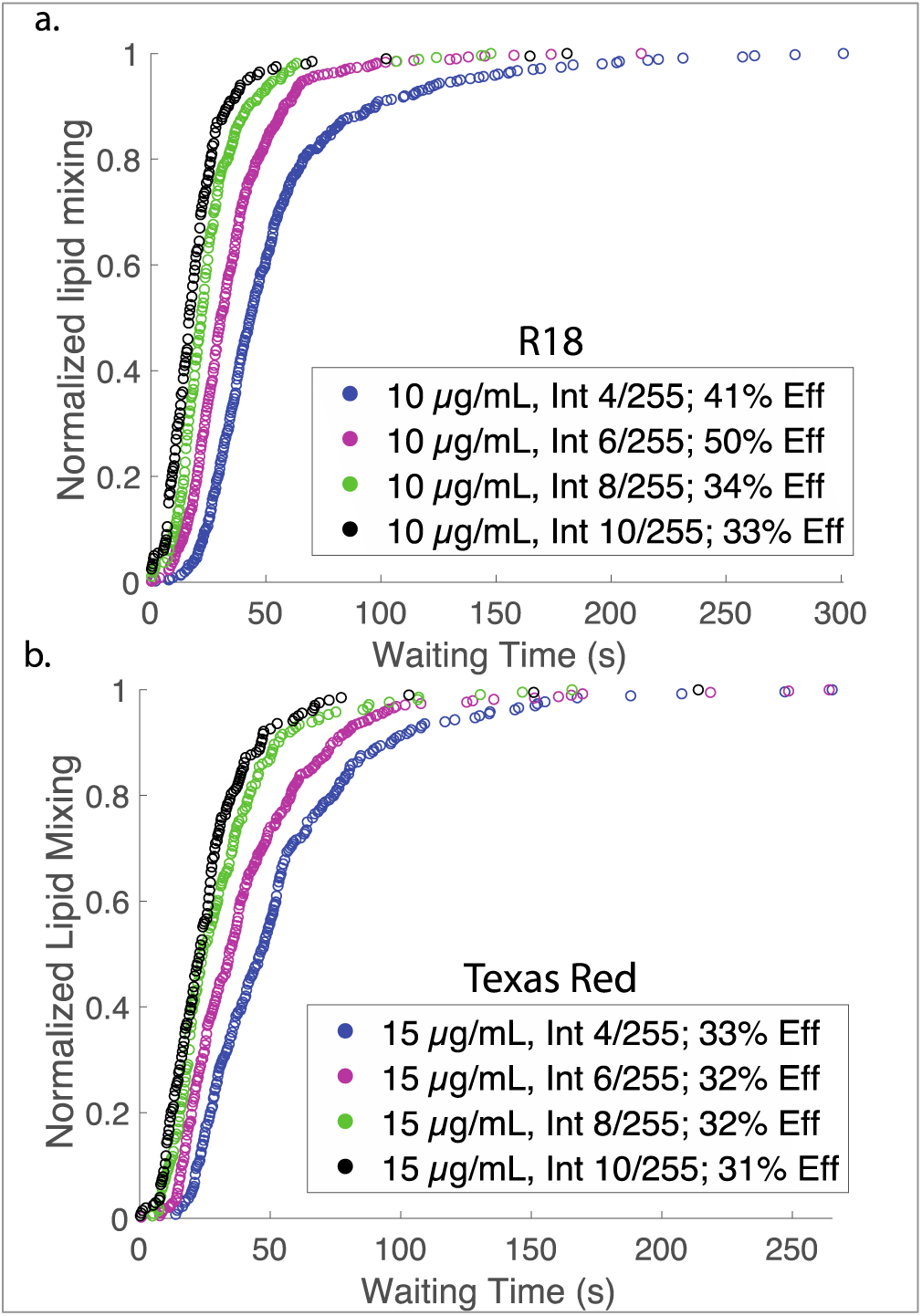
Kinetics of viral hemifusion perturbed by higher light intensity. Cumulative distribution plots for lipid mixing of individual viruses labeled with R18 (a) or Texas Red (b) are shown at different illumination intensities. Legends denote the dye concentration used for labeling, the intensity of illumination relative to the max LED output of 255, and the efficiency of lipid mixing (denoted Eff). Higher illumination leads to faster lipid-mixing kinetics.

Dyes conjugated to two-tailed lipids are more difficult to load into viral particles at self-quenching concentration, but several, including Texas Red-DHPE, have been used as probes for membrane fusion in different viral systems (3, 22-25). We have anecdotally found Texas Red more robust to photoinhibition effects (14, 15), so we tested it more systematically. At the lowest concentrations tested, Texas Red was more robust to photoinhibition effects than R18, although photoinhibition was evident at higher concentrations and illumination intensities. The mean numbers of dyes per virion are listed in Table 1 for each labeling concentration used; fluorescence dequenching ratios are given in Fig. S1. An increase in the rate of fusion kinetics was evident with increasing illumination, demonstrating that fusion can be perturbed even in the absence of photoinactivation. Here, we use photoinhibition to refer broadly to reduction of lipid mixing and subsequent fusion after a given waiting time and photoinactivation to refer more specifically to the reduction of the maximum amount of virus undergoing fusion, presumably in an irreversible fashion. Cumulative distribution functions for influenza fusion using Texas Red-labeled and R18-labeled virus overlaid well at the lowest dye concentrations and illumination intensities used, and no significant photoinactivation was detected. This asymptotic behavior suggests that both of these dyes can yield accurate measures of viral membrane fusion if dye concentration and illumination intensity are tightly controlled.

**Table 1:** Labeling efficiency of influenza virus with R18 or Texas Red. At low dye concentrations, there is a supra-linear increase in labeling with dye concentration; this becomes linear in the 20-35 µg/mL region for both dyes and begins to saturate at 40 µg/mL for R18.

| **R18**<br>**Labeling solution** | **R18 mean**<br>**dyes per virion** | **Texas Red**<br>**Labeling solution** | **Texas Red mean**<br>**dyes per virion** |
| --- | --- | --- | --- |
| 10 µg/mL | 1900 | 15 µg/mL | 940 |
| 20 µg/mL | 5100 | 25 µg/mL | 5500 |
| 30 µg/mL | 6800 | 35 µg/mL | 8000 |
| 40 µg/mL | 7100 |  |  |

To probe the chemical nature of the photoinactivation process, we hypothesized that dyes might form photoadducts with viral components. We tested this for viral proteins by adding dye to viral samples, either exposing them to light or not, and then detergent-solubilizing the samples and size-separating proteins *via* gel electrophoresis. As shown in Fig. 4, viral envelope or membrane-proximal proteins acquired red fluorescence after illumination in a dye-concentration-dependent manner in R18-labeled samples but not detectably in Texas Red-labeled samples. In particular, viral hemagglutinin undergoes a gel shift upon R18 labeling and illumination but not upon Texas Red labeling and illumination; the fluorescence spectrum of the shifted band is shown in Fig. S3. This demonstrates a covalent adduct formed by R18 dye and a portion of the hemagglutinin. These results suggest that R18:viral protein photoadducts correlate with photoinactivation, although they do not prove that the protein adducts, as opposed to other photochemical products in the viral membrane, are causative for viral inactivation.

**Figure 4:**
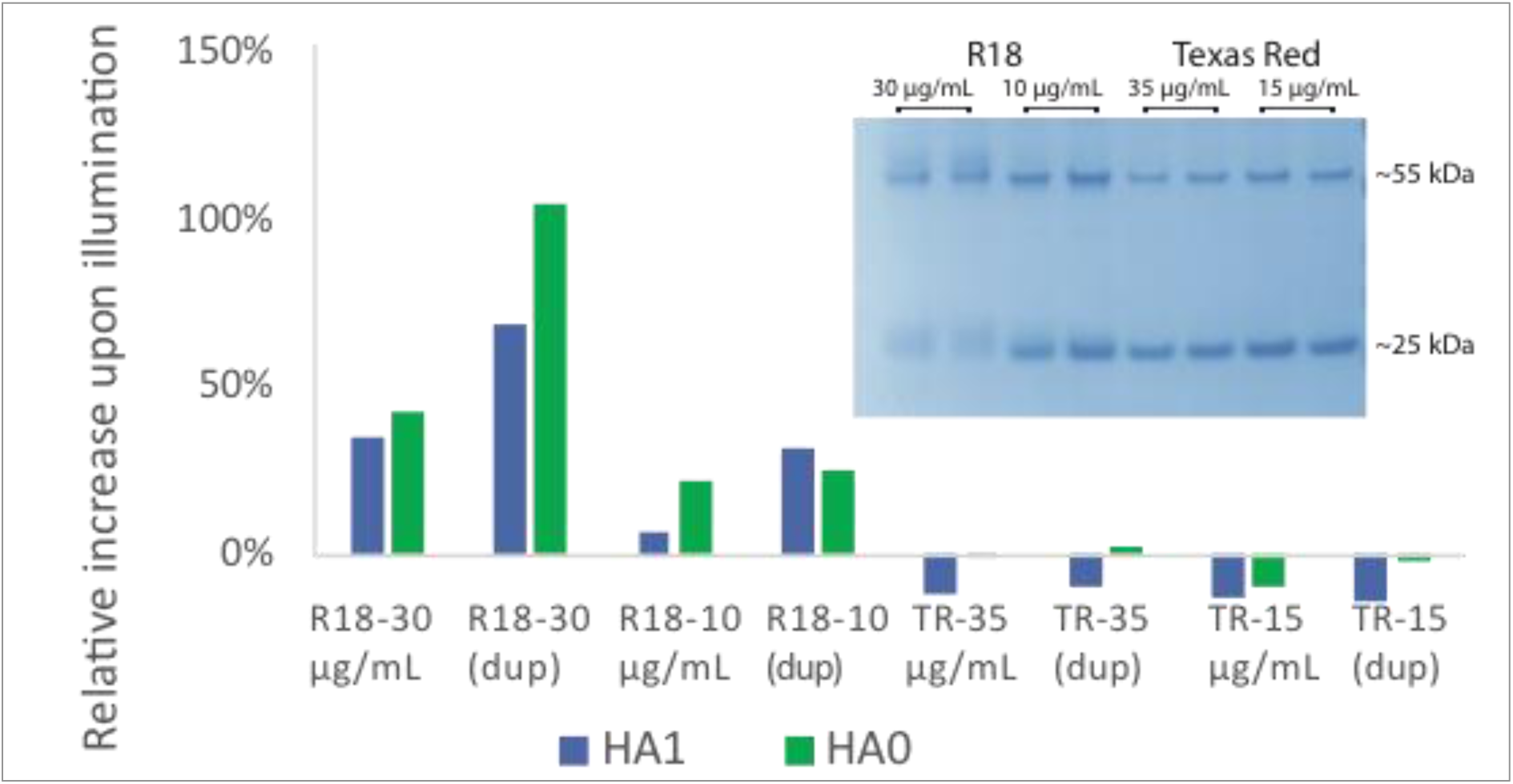
R18 forms photoadducts with influenza proteins upon illumination. SDS-PAGE of dye-labeled influenza virus with and without illumination yielded fluorescent protein bands. Fluorescence intensities of the bands corresponding to the HA1 and HA0 subunits (∼55 kDa and ∼75 kDa, respectively) were calculated, these were normalized by the Coomassie Blue staining intensity of each band, and then the % increase in fluorescence was calculated between unilluminated and illuminated samples. Samples are denoted by the dye (R18 or Texas Red) and concentration (µg/mL) used to label the virus; “dup” denotes duplicate samples. The use of R18 as a lipid marker caused a substantial increase in protein fluorescence in a dye-concentration-dependent manner, while Texas Red-DHPE did not when similarly introduced into the viral membrane. A gel shift is for the 25-kDa band corresponding to HA2 and the 55-kDa band corresponding to HA1 is also evident in the Coomassie Blue-stained bands after illumination of the R18-labeled samples, as shown in the inset. Full gels are shown in Fig. S4.

To test whether photoinactivation can explain the observed dye and illumination effects on fusion, we used two sets of kinetic models, one simple chemical kinetics model similar to that previously used to analyze Zika virus inactivation (14) but modified for influenza (see Methods) and one cellular-automaton model that can explicitly treat inactivation of individual hemagglutinin trimers (19-21). As schematized in Figure 5, the cellular automaton model describes state transitions of individual hemagglutinin trimers (“per-protein model”) whereas the chemical kinetics model is formulated to describe state transitions of whole virus particles (“per-virus model”). Fusion and inactivation parameters for each of these models were optimized to fit fusion efficiencies and waiting-time distributions (see Methods for details). Both models produced a decrease in fusion efficiency with increasing R18 concentration or illumination (Fig. 5), but neither could well explain the increase in fusion rates observed. Interestingly, the cellular-automaton model fit the efficiency data much better over the full range of dye concentrations and illumination intensities measured. Since in the chemical kinetics model the whole virus inactivates as a unit, while in the cellular automaton model each hemagglutinin trimer undergoes stochastic inactivation independently, the better fit of the cellular automaton model suggests that dye photoreaction with individual fusion proteins better describes the physical process of photoinactivation. These results are consistent with the following explanation: reduced fusion efficiency with increased R18 dye concentration or illumination power results from hemagglutinin inactivation due to photoconjugation. Interestingly, a previous report noted photoinactivation of Sindbis virus by R18 without inhibition of lipid mixing, in that case without detectable photoreactivity on SDS-PAGE (26). In our findings, the increased rates of lipid-mixing likely reflect another mechanistic process, possibly dye-mediated photoconjugation or photo-oxidation of viral lipids or alternately a local heating effect from dye thermal relaxation.

**Figure 5:**
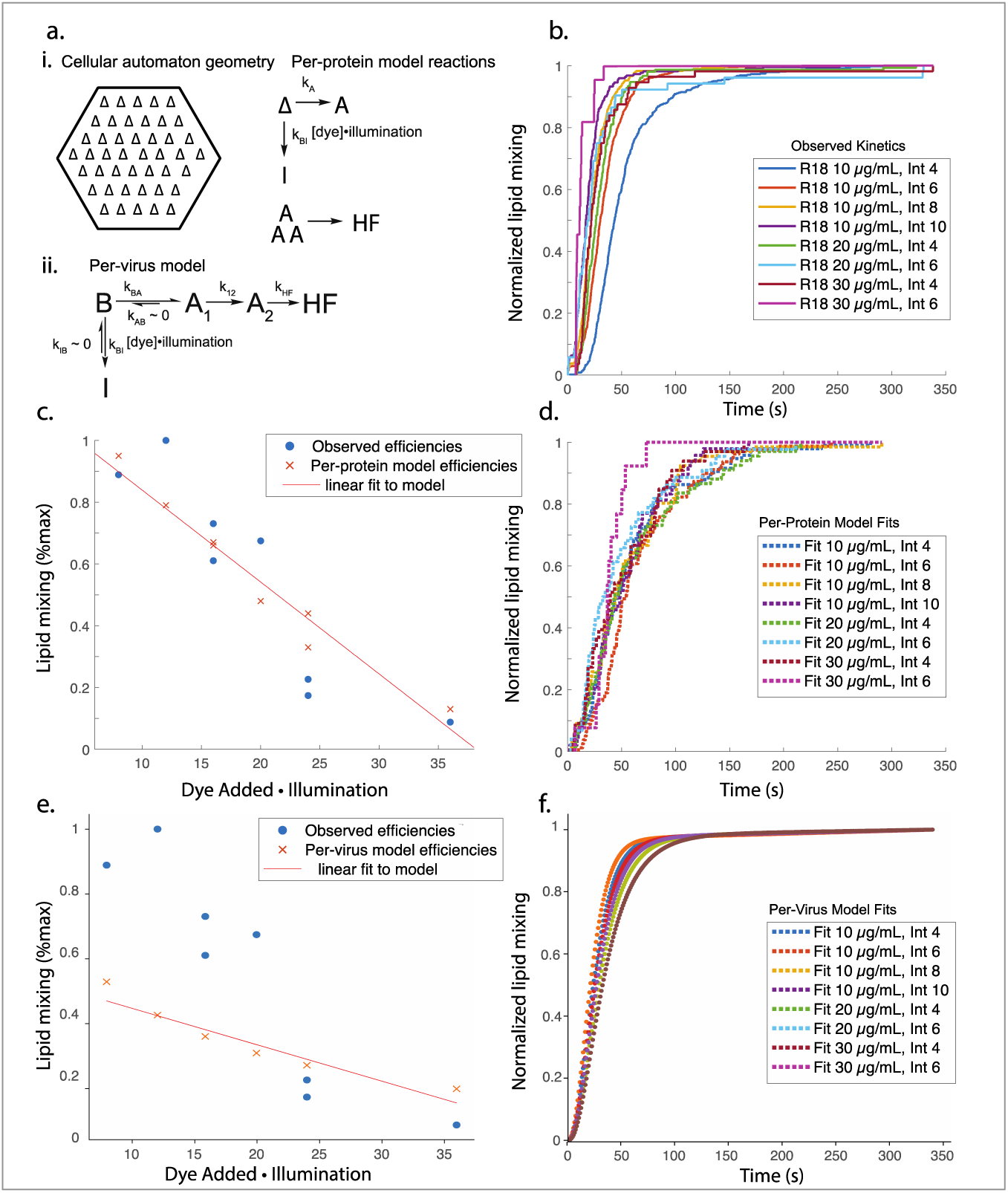
Kinetic models of fusion protein inactivation can account for photoinactivation of fusion. Computational models of per-fusion-protein inactivation and per-virus inactivation (schematized in panel a) were fit to single-virus fusion data at multiple R18 dye concentrations and illumination intensities (cumulative distribution functions in panel b). The models specify interconversion rates among bound (B), activated (A), inactivated (I), and hemifused (HF) states. In the per-protein model, states describe individual hemagglutinin trimers (except HF), whereas in the per-virus model, states describe the virus as a whole. The per-protein model could better account for the reduced fusion efficiency than the per-virus model (panels c, e) when lipid mixing was normalized across conditions. Cumulative distribution functions of modeled and observed fusion events normalized within each dye and illumination condition (panels b, d, f) show that neither model robustly reproduces the slight speeding of fusion at increased dye.illumination. Total numbers of lipid-mixing events used to construct the cumulative distribution events were 482, 677, 294, 222, 153, 52, 56, and 22 for the eight conditions plotted. The cellular automaton used in the model represents the virus:target membrane contact zone as a hexagonal lattice with individual hemagglutinin trimers that can convert to activated (A) or inactive (I). Three neighboring activated trimers result in hemifusion. Legend in panels b, d, and f specifies dye loading concentration, and intensity (out of 255).

## Conclusions

Our data demonstrate that photoreactivity of common fluorescent dyes can inhibit viral membrane fusion and perturb apparent fusion kinetics. Careful optimization of dye labeling and illumination intensity can avoid measurable photoreactivity and perturbation, but care should be taken in designing and interpreting membrane fusion experiments to avoid results that reflect photoreactivity and photoinhibition, and in order to make meaningful comparisons among results from different laboratories. In the case of influenza membrane fusion, R18 is particularly prone to a photoinhibition effect that can be largely explained by photoconjugation of the dye to the hemagglutinin protein. Both R18 and Texas Red perturb fusion kinetics in an illumination-dependent manner that cannot be explained by photoconjugation. These can be avoided using low dye concentrations and minimal illumination. Furthermore, such effects are thus likely to be quite general and extend beyond influenza viral fusion and even viral membrane fusion, although these systems provide sensitive readouts of dye photoreactivity. Fluorescent probes provide powerful chemical tools to measure biochemical processes, but the possibility to perturb the system of interest is always a concern, and these results provide concrete data of how viral membrane fusion can be perturbed by dye photoreactivity. These results also argue for assay systems that minimize dye levels in the virions and avoid the light-dependent processes documented here.

## Author Contributions

R.J.R designed experiments, performed experiments, analyzed data, and co-wrote the paper. A.M.V.G. performed experiments and analyzed data. S.G.B. designed experiments and co-wrote the paper. P.M.K designed experiments, analyzed data, and co-wrote the paper.

## Acknowledgements

The authors would like to thank K. Liu, K. Okamoto, A. Sengar, and E. Webster for helpful discussions. This work was supported by a Wallenberg Academy Fellowship to P.M.K. and National Institutes of Health grants R35 GM118044 to S.G.B and R01 GM098304 to P.M.K.

## References

1. Blumenthal, R., S. A. Gallo, M. Viard, Y. Raviv, and A. Puri. 2002. Fluorescent lipid probes in the study of viral membrane fusion. Chem Phys Lipids 116(1-2):39–55.

2. Morris, S. J., D. P. Sarkar, J. M. White, and R. Blumenthal. 1989. Kinetics of pH-dependent fusion between 3T3 fibroblasts expressing influenza hemagglutinin and red blood cells. Measurement by dequenching of fluorescence. J Biol Chem 264(7):3972–3978.

3. Razinkov, V. I., G. B. Melikyan, R. M. Epand, R. F. Epand, and F. S. Cohen. 1998. Effects of spontaneous bilayer curvature on influenza virus-mediated fusion pores. J Gen Physiol 112(4):409–422.

4. Schroth-Diez, B., E. Ponimaskin, H. Reverey, M. F. Schmidt, and A. Herrmann. 1998. Fusion activity of transmembrane and cytoplasmic domain chimeras of the influenza virus glycoprotein hemagglutinin. J Virol 72(1):133–141.

5. Blumenthal, R., A. Balipuri, A. Walter, D. Covell, and O. Eidelman. 1987. Ph-Dependent Fusion of Vesicular Stomatitis-Virus with Vero Cells - Measurement by Dequenching of Octadecyl Rhodamine Fluorescence. Journal of Biological Chemistry 262(28):13614–13619.

6. Stegmann, T., H. W. Morselt, J. Scholma, and J. Wilschut. 1987. Fusion of influenza virus in an intracellular acidic compartment measured by fluorescence dequenching. Biochim Biophys Acta 904(1):165–170.

7. MacDonald, R. I. 1985. Membrane fusion due to dehydration by polyethylene glycol, dextran, or sucrose. Biochemistry 24(15):4058–4066.

8. Hughes, L. D., R. J. Rawle, and S. G. Boxer. 2014. Choose your label wisely: water-soluble fluorophores often interact with lipid bilayers. PLoS One 9(2):e87649.

9. Nunes-Correia, I., A. Eulalio, S. Nir, N. Duzgunes, J. Ramalho-Santos, and M. C. Pedroso de Lima. 2002. Fluorescent probes for monitoring virus fusion kinetics: comparative evaluation of reliability. Biochim Biophys Acta 1561(1):65–75.

10. Bowen, M. E., K. Weninger, A. T. Brunger, and S. Chu. 2004. Single molecule observation of liposome-bilayer fusion thermally induced by soluble N-ethyl maleimide sensitive-factor attachment protein receptors (SNAREs). Biophys J 87(5):3569–3584.

11. Floyd, D., J. R. Ragains, J. J. Skehel, S. C. Harrison, and A. M. van Oijen. 2008. Single-particle kinetics of influenza virus membrane fusion. Proceedings of the National Academy of Sciences 105(40):15382–15387.

12. Wessels, L., M. W. Elting, D. Scimeca, and K. Weninger. 2007. Rapid membrane fusion of individual virus particles with supported lipid bilayers. Biophys J 93(2):526–538.

13. Schmidt, F. I., P. Kuhn, T. Robinson, J. Mercer, and P. S. Dittrich. 2013. Single-virus fusion experiments reveal proton influx into vaccinia virions and hemifusion lag times. Biophys J 105(2):420–431.

14. Rawle, R. J., E. R. Webster, M. Jelen, P. M. Kasson, and S. G. Boxer. 2018. pH dependence of Zika membrane fusion kinetics reveals an off-pathway state. ACS Central Science.

15. Rawle, R. J., S. G. Boxer, and P. M. Kasson. 2016. Disentangling Viral Membrane Fusion from Receptor Binding Using Synthetic DNA-Lipid Conjugates. Biophys J 111(1):123–131.

16. Hughes, L. D. 2013. Model membrane architectures for the study of membrane proteins.

17. Floyd, D. L., S. C. Harrison, and A. M. van Oijen. 2009. Method for measurement of viral fusion kinetics at the single particle level. JoVE (Journal of Visualized Experiments)(31):e1484.

18. Hutchinson, E. C., P. D. Charles, S. S. Hester, B. Thomas, D. Trudgian, M. Martinez-Alonso, and E. Fodor. 2014. Conserved and host-specific features of influenza virion architecture. Nature communications 5:4816.

19. Ivanovic, T., J. L. Choi, S. P. Whelan, A. M. van Oijen, and S. C. Harrison. 2013. Influenza-virus membrane fusion by cooperative fold-back of stochastically induced hemagglutinin intermediates. Elife 2:e00333.

20. Ivanovic, T., and S. C. Harrison. 2015. Distinct functional determinants of influenza hemagglutinin-mediated membrane fusion. Elife 4.

21. Zawada, K. E., K. Okamoto, and P. M. Kasson. 2018. Influenza Hemifusion Phenotype Depends on Membrane Context: Differences in Cell-Cell and Virus-Cell Fusion. J Mol Biol 430(5):594–601.

22. Tse, F. W., A. Iwata, and W. Almers. 1993. Membrane flux through the pore formed by a fusogenic viral envelope protein during cell fusion. J Cell Biol 121(3):543–552.

23. Lakadamyali, M., M. J. Rust, H. P. Babcock, and X. Zhuang. 2003. Visualizing infection of individual influenza viruses. Proc Natl Acad Sci U S A 100(16):9280–9285.

24. van der Schaar, H. M., M. J. Rust, C. Chen, H. van der Ende-Metselaar, J. Wilschut, X. Zhuang, and J. M. Smit. 2008. Dissecting the cell entry pathway of dengue virus by single-particle tracking in living cells. PLoS Pathog 4(12):e1000244.

25. Zaitseva, E., S. T. Yang, K. Melikov, S. Pourmal, and L. V. Chernomordik. 2010. Dengue Virus Ensures Its Fusion in Late Endosomes Using Compartment-Specific Lipids. PLoS Pathogens6(10).

26. Thongthai, W., and K. Weninger. 2009. Photoinactivation of sindbis virus infectivity without inhibition of membrane fusion. Photochem Photobiol 85(3):801–806.

